# Fatigue induces long lasting detrimental changes in motor skill learning

**DOI:** 10.1101/406520

**Authors:** Meret Branscheidt, Panagiotis Kassavetis, Davis Rogers, Martin A. Lindquist, Pablo Celnik

**Author notes:** Corresponding author: Pablo Celnik, MD 600 N. Wolfe St. Baltimore, MD 21287, USA and Meret Branscheidt, MD.

## Abstract

Fatigue due to physical exertion is a ubiquitous phenomenon in everyday life and especially common in a range of neurological diseases. While the effect of fatigue on limiting skill execution are well known, its influence on learning new skills is unclear. This is of particular interest as it is common practice to train athletes, musicians or perform rehabilitation exercises up to and beyond a point of fatigue. In a series of experiments, we describe how fatigue impairs motor skill learning beyond its effects on task execution. The negative effects on learning are evidenced by impaired task acquisition on subsequent practice days even in the absence of fatigue. Further, we found that this effect is in part mediated centrally and can be alleviated by altering motor cortex function. Thus, the common practice of training while, or beyond, fatigue levels should be carefully reconsidered, since this affects overall long-term skill learning.

## INTRODUCTION

We know from everyday life that, in order to gain and maintain proficiency, the single most important requirement in a motor skill is practice. Intensive, repetitive training is an essential routine for musicians, artists, surgeons, and athletes. Repetitive practice is also part of rehabilitation approaches to recover function of the motor system and other domains. While repetition improves performance over time, there comes a point when it also causes fatigue and an eventual degradation of task execution (Boyas & Guével, 2011; Gandevia, Enoka, McComas, Stuart, & Thomas, 1995b).

Studies investigating fatigue have made a distinction between fatigue as a cognitive phenomenon and fatigue as a neuromuscular phenomenon (Janet L, 2012), although this separation can be blurred at times (Kuppuswamy, 2017). In neurological conditions, for instance, fatigue has been described as an overall state linked to changes in motor cortex excitability (Kuppuswamy, Clark, Turner, Rothwell, & Ward, 2015). Here, we use the term fatigue to describe the degradation of maximal force output induced through physical exertion of task relevant muscles that is associated with a perceptual state with heightened effort.

Surprisingly little is known about the effects of muscle fatigue on the acquisition of motor skills. The existing literature regarding motor learning under fatigue is mostly limited to a few studies from the 1970-90s with contradictory results (for a comprehensive overview see (Janet L, 2012). While some studies have reported that participants are unable to learn a motor task under fatigue conditions (Carron and Ferchuk, 1971, Thomas et al., 1975), others have not found fatigue to be detrimental to motor learning (Cotten et al., 1972, Alderman, 1965, Spano and Burke, 1976). One key challenge in studying motor learning under fatigue is the so-termed “performance-learning” distinction (Cahill, McGaugh, & Weinberger, 2001; Kantak & Winstein, 2012): Performance is usually defined as a temporary effect; e.g. how skillful a movement is executed during one training session. In contrast, learning can only be inferred indirectly from performance, via measuring differences in performance over time or over tasks (Kantak & Winstein, 2012). This distinction is important because experimental conditions that affect performance do not necessarily have to affect learning. For example, while the performance of rats in the absence of a motivational cue seemed to show no learning in a maze task despite repeated practice, providing a food reward uncovered that they indeed were able to learn the right path nonetheless (Tolman & Honzik, 1930). Thus, it is necessary to separate decreased task performance under fatigue with true effects of fatigue on motor learning. Here, we address this issue by disentangling the effect of muscle fatigue on learning a motor skill from the performance confounder.

In experiment 1 (*N* =38), we asked healthy individuals to learn a sequential pinch force task over two days and showed that, even though participants were only fatigued at Day 1, skill learning was impaired on both days. Interestingly, a subgroup of fatigued subjects (*N* =12) took two additional days of training with no fatigue to catch up to the skill performance level of the non-fatigued group. In experiment 2 (*N* =20), we tested performance on the untrained, unfatigued hand and demonstrated that participants have impaired skill learning in both the fatigued and unfatigued effector. Finally, in experiment 3 (*N* =45), we replicated the findings of experiment 1 and tested whether the negative effects of fatigue on learning are centrally mediated. We found that disruptive rTMS to the motor cortex (Cantarero, Lloyd, & Celnik, 2013a; Huang et al., 2010) partly alleviates the adverse effects of fatigue on skill learning, suggesting a possible role for maladaptive memory formation under fatigued conditions.

Altogether, the results of the three experiments provide the first evidence that fatigue has a lasting adverse effect on motor skill learning beyond performance. The findings are highly significant to all professions that rely on intensive physical training to achieve optimal performance. Understanding the effects of fatigue on learning helps the formulation of training and rehabilitation regimens geared to improve motor function. Although not explored here, similar effects might be present in other learning domains (i.e. in cognition, perception).

## RESULTS

### Fatigue has lasting effects on acquisition of motor skill

In the first experiment, we assessed how muscle fatigue influenced skill learning over multiple days. 38 participants trained in an isometric pinch task over the course of two days; see Fig 1. While all subjects were instructed to perform an isometric pinch contraction prior to four bouts of training, a subset of participants (Fatigue group (FTG), *N* =20) performed the contractions until experiencing muscle fatigue (∼60% decrement of maximal voluntary contraction, MVC measured in Newton and monitored by surface electromyographic (EMG) signal) on Day 1. On Day 2 both groups performed the skill task without the induction of fatigue. Skill learning was indexed by a measure that quantifies shifts in the relationship between movement time and accuracy rate (Reis et al., 2009). As the relationship between learning rate and skill measure appeared linear, a regression line was fit separately for each day and group. Here the slope of the regression line represents the learning rate (see Methods).

**Figure 1.**
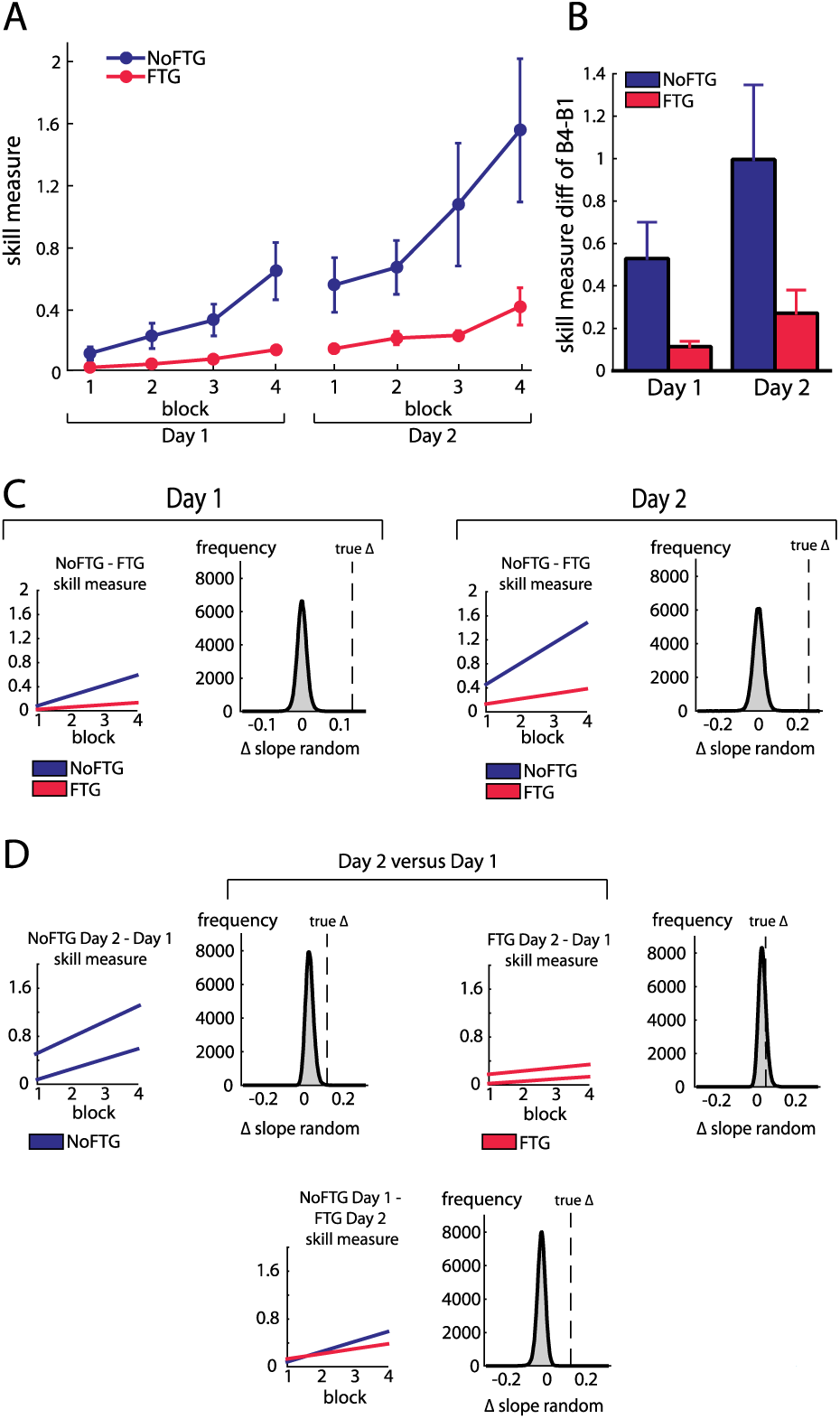
Comparison of skill acquisition in an isometric pinch task between fatigued and non-fatigued participants. Panel (A) shows changes in skill measure over the course of four training blocks on two consecutive days for both groups (NoFTG =blue; FTG =red). While both groups improved task execution, the FTG had a lower performance level on Day 1 and on Day 2 when they were not fatigued compared to controls. Note that the skill performance in block 4 of Day 2 in the FTG remained below the level of NoFTG at the end of Day 1. Panel (B) shows the difference in performance between block 4 to block 1 for both groups on each day. Panel (C) shows the differences in learning rates for Day 1 and Day 2 between groups. We compared the learning rate of both groups by first fitting a robust linear regression model to the individual data of each group (line plots). To test if there was a true difference in learning rates, we calculated the Δ of the average slope of the fitted model (regression coefficients). We then used permutation testing to generate a distribution of Δ slopes of randomly generated groups (grey histoplots). The Null hypothesis was rejected when less then 5% of the generated Δ slopes exceed the true Δ slope (dashed vertical lines, see also methods). In Panel (D) learning rates are compared within groups across days (Day 2 versus Day 1). There were no significant differences in learning rates across days in both groups.

Fatigue was reliably induced during Day 1 in the fatigue group, as shown by decrements of MVC over time. Importantly, MVC always stayed above the force level required to execute the task (up to 40% of MVC; see Supplementary Information). All participants improved their ability to execute the task on both days; see Fig 1. However, on both days, learning rates for the non-fatigue group (NoFTG) were significantly larger than for the FTG (Day 1: mean slopes NoFTG 0.169 versus FTG 0.038; p =0.01. Day 2: mean slopes NoFTG 0.339 versus FTG 0.083; p =0.03; Fig 1). It is important to note that the lower performance of the fatigued group on Day 1 does not allow to make a direct inference about lower motor learning (f. e. because of task differences). Of note, even on Day 2 when the FTG was not fatigued and both groups were performing the same task, the ability to execute the task did not reach the level of the NoFTG subjects on Day 1 despite having twice the amount of practice (p =0.01). There were no changes in overall learning rate across days for either controls or the fatigued group (Day 1 versus Day 2 NoFTG: p =0.21, FTG: p =0.77), indicating that learning rates remained low in the FTG.

A separate analysis of the changes in movement time and in percentage of correct trials showed that the effect of fatigue on skill performance was due to more errors in the FTG and not to differences in movement time. Counter intuitively, this higher rate of errors was due to increased force production in the FTG resulting in target overshoot. This was most prominent for the two lower force targets while performance in the highest force target was similar between groups (see also Supplementary Information).

To assess, how long the FTG took to reach similar performance levels as the NoFTG, a subgroup of participants continued training up to four days (NoFTG_4D_ *N* =12, FTG_4D_ *N* =12). While Day 1 & 2 showed the same result with lower learning rates for the FTG_4D_ (Day 1: p < 0.01; Day 2: p < 0.01), this group reached similar performance levels to the control group only towards the end of Day 3 and on Day 4 (Day 3: p =0.07; Day 4: p =0.09; see Fig 2).

**Figure 2.**
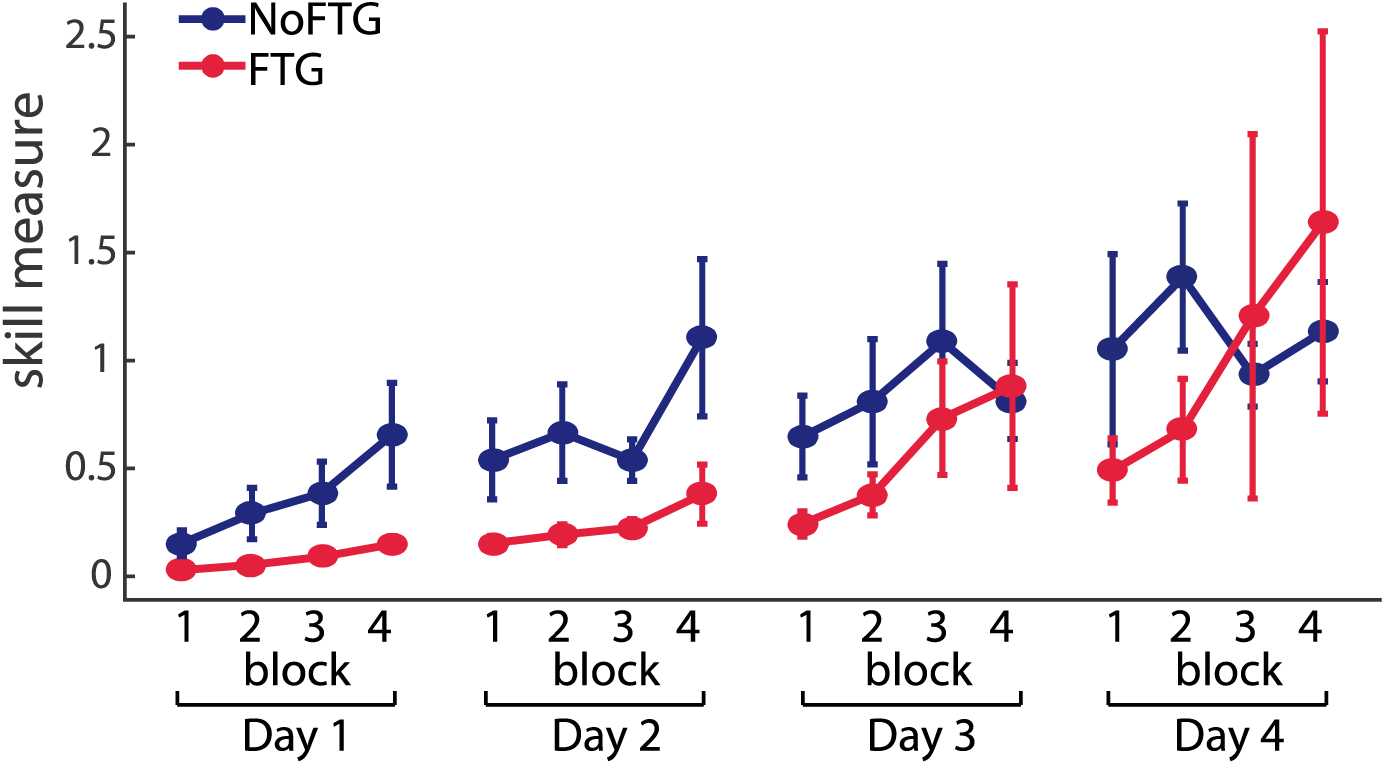
Comparison of skill execution between FTG and NoFTG over the course of four days. Note that the FTG (red) showed lower skill levels at Day 1 & 2 compared to the NoFTG (blue) and only reached similar levels to controls at the end of Day 3 and on Day 4.

Since learning in the FTG was impaired even in the non-fatigued state on Day 2, these results indicate that learning under fatigue conditions has an adverse long-lasting effect on skill acquisition.

### Fatigue affects performance even in the untrained, non-fatigued effector

Because execution under fatigue is impaired, it is conceivable that this performance confounder masked skill learning. Assessing the transfer of learning to the uninstructed, unfatigued hand provides a unique way to circumvent this challenge. Generalization of motor skills across hands has previously been well characterized, where skill training with one hand results in improved performance in the untrained hand (Camus, Ragert, Vandermeeren, & Cohen, 2009; Perez, Wise, Willingham, & Cohen, 2007). Thus, in experiment 2, we measured skill execution in the left hand of a new group of 20 participants before and after training with their right fatigued (FTG_TRANSFER_, *N* =10) or non-fatigued hand (NoFTG_TRANSFER_, *N* =10).

Similar to experiment 1, right hand learning rates over the four blocks were lower in participants that performed the task under fatigue compared to controls (mean slope NoFTG_TRANSFER_ 0.03 versus FTG_TRANSFER_ 0.008; p =0.01; Fig 3). As expected, prior to training, the skill measure of the left hand was similar between groups (t_18_ =−0.157, p =0.88). After training with the right hand, performance of the left hand was significantly lower in the FTG_TRANSFER_ compared to the NoFTG_TRANSFER_ (mean Δblock2-block1 NoFTG_TRANSFER_ 0.133 ±0.036 versus FTG_TRANSFER_ 0.018 ±0.003; p < 0.01; Fig 3).

**Figure 3.**
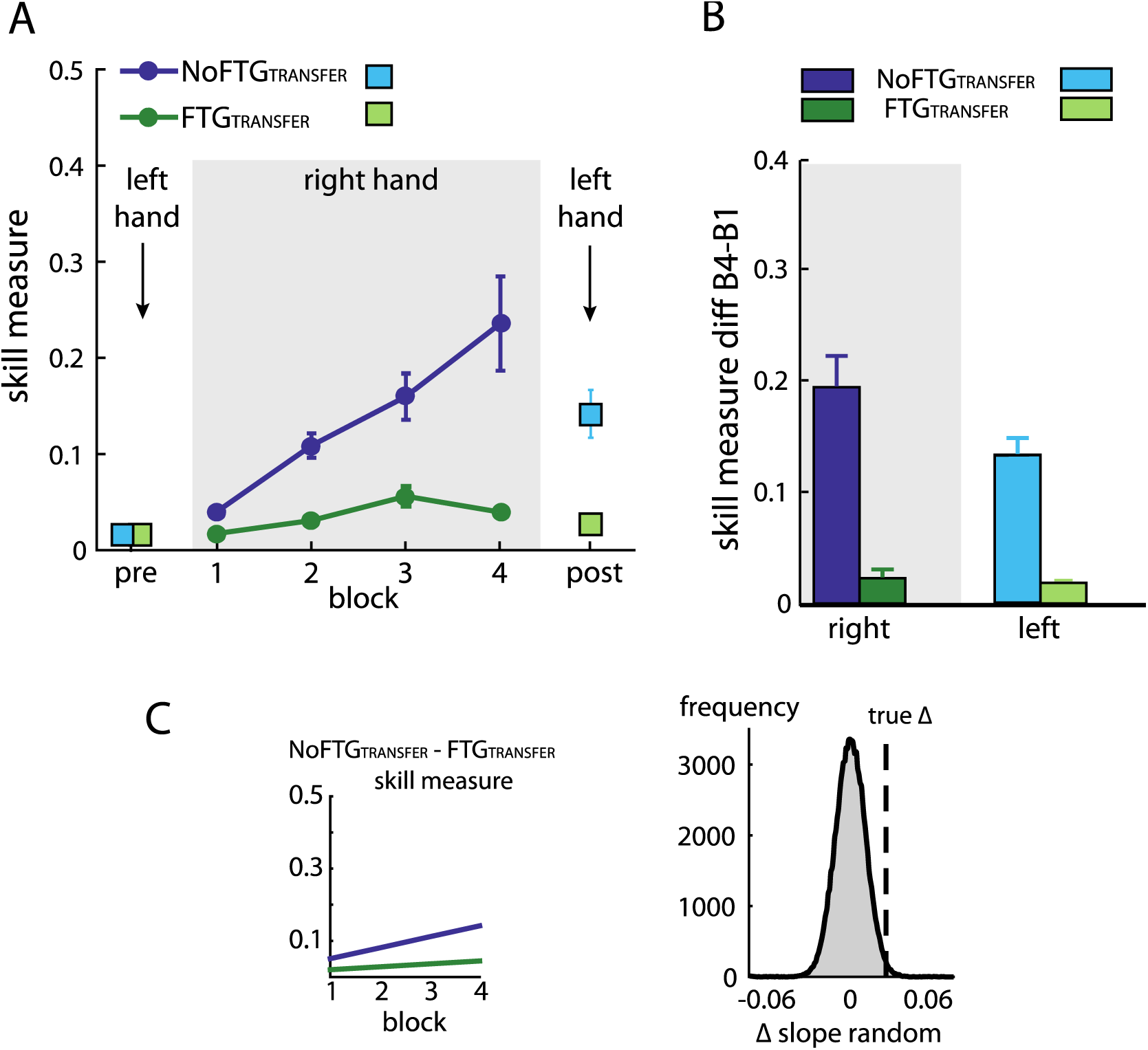
Intermanual transfer of learning in fatigued versus non-fatigued participants. Panel (A) shows changes in the skill measure over the course of four blocks during one day of training (NoFTG_TRANSFER_ =dark blue, FTG_TRANSFER_ =dark green). Before and after the training, both groups performed 15 trials of the pinch force task with their left hand (FTG_TRANSFER_ =light blue square, NoFTG_TRANSFER_ =light green square). Note that, while both groups improved skill performance over time, the FTG_TRANSFER_ group had a lower performance level, consistent with experiment 1, in both the fatigued and non-fatigued effector. Panel (B) shows the difference in performance between block 4 to block 1 for the right hand and block 2 - block 1 for the left hand in both groups. Performance in the left hand was significantly lower in the FTG_TRANSFER_ compared to NoFTG_TRANSFER_. Panel (C) shows the learning rates and the true Δ slope in comparison to randomly generated Δ slopes after permutation. Similar to experiment 1, controls showed higher learning rates than the fatigued group.

Together, these results indicate that fatigue affected learning and that the performance confounder did not mask the expression of learning. Importantly, the poor performance in the untrained hand suggests that fatigue impairs central motor skill learning mechanisms beyond any potential adverse effect in the fatigued effector.

### Long lasting detrimental effects of fatigue on learning are centrally mediated

To determine whether the effect of fatigue on learning is centrally mediated, we interfered with primary motor cortex processes thought to be involved in skill retention in a new group of participants (Galea, Vazquez, Pasricha, de Xivry, & Celnik, 2011; Muellbacher et al., 2002; Reis et al., 2009; Richardson et al., 2006). To this end, in experiment 3 we used disruptive rTMS (repetitive transcranial magnetic stimulation; (Cantarero et al., 2013a; Huang et al., 2010) over the primary motor cortex (M1) after task training on Day 1 (FTG_M1_, *N* =15). To control for potentially non-specific effects of rTMS, we also tested a fatigued and a non-fatigued group with TMS applied over the parietal interhemispheric fissure (Pz, according to 10-20 system; FTG_SHAM_, *N* =15 and NoFTG_SHAM_, *N* =10).

The permutation test showed that learning rates of both fatigued groups were smaller compared to controls, but similar to each other on Day 1 (mean slope NoFTG_SHAM_ 0.049, FTG_SHAM_ 0.02, FTG_M1_ 0.016; NoFTG_SHAM_ versus FTG_SHAM_ p =0.01, NoFTG_SHAM_ versus FTG_M1_ p < 0.01, FTG_SHAM_ versus FTG_M1_ p =0.73; Fig 4). On Day 2, consistent with experiment 1 and 2, the learning rate was still smaller in FTG_SHAM_ compared to the NoFTG_SHAM_ control. However, the learning rate of the FTG_M1_ group was *not* significantly different from the NoFTG_SHAM_, but significantly different from FTG_SHAM_ (mean slope NoFTG_SHAM_ 0.04, FTG_SHAM_ 0.022, FTG_M1_ 0.042; NoFTG_SHAM_ versus FTG_SHAM_ p =0.04, NoFTG_SHAM_ versus FTG_M1_ p =0.47, FTG_SHAM_ versus FTG_M1_ p =0.03). Of note, comparing learning rates across days within groups, we found a significant difference for FTG_M1_ (Day 1 versus Day 2, FTG_M1_ p =0.04), but no difference for the other two groups (Day 1 versus Day 2, NoFTG_SHAM_ p =0.94, FTG_SHAM_ p =0.88).

**Figure 4:**
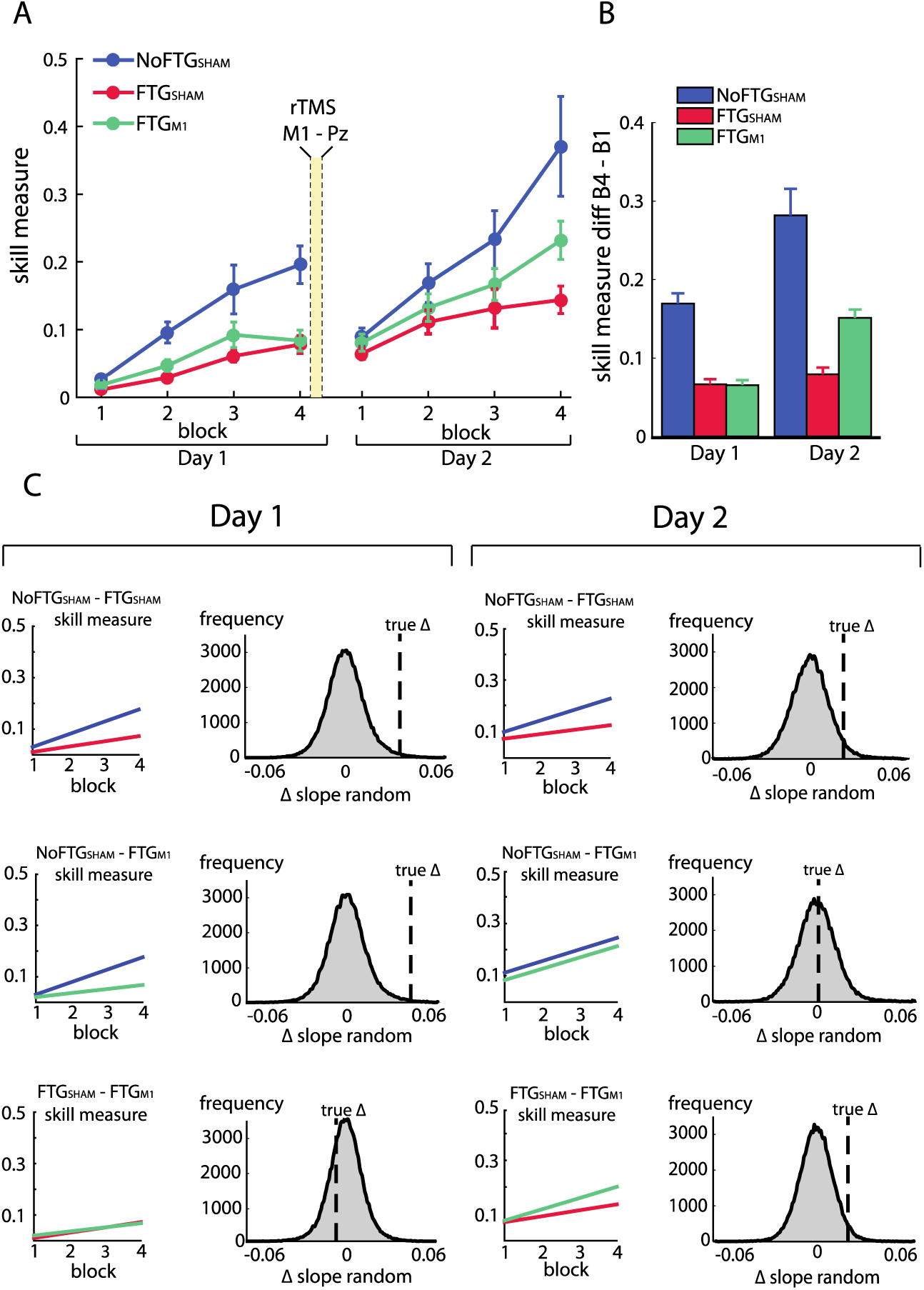
Disrupting M1 at the end of Day 1 training reduces the impaired skill acquisition on Day 2. Panel (A) shows changes in the skill measure over four training blocks on two consecutive days for all groups (NoFTG_SHAM_ =blue, FTG_SHAM_ =red, FTG_M1_ =green). Note that the FTG_M1_ experienced higher improvement of performance in Day 2 compared to the FTG_SHAM_. Panel (B) shows the difference in performance between block 4 to block 1 for all groups on each day. Panel (C) shows the comparison of learning rates for all groups on Day 1 and 2. On both days, the unfatigued control group showed a higher learning rate than the FTG_SHAM_ group (this group received sham stimulation at the end of Day 1). Second row: In contrast, there was no significant difference in learning rates between the FTG_M1_ group and the control group on Day 2 (in this group, M1 function was disrupted at the end of Day 1 using rTMS), while learning rates on Day 1 were lower. Third row: Both groups showed similar performance on Day 1, but higher learning rates were evidenced on Day 2 in the FTG_M1_ group compared to the FTG_SHAM_ group.

Together, these results show that disruption of M1 function after training diminished the detrimental effects of fatigue on motor skill learning. This indicates that the long-lasting effects of fatigue on learning are at least partly centrally mediated and linked to motor memory formation.

### Fatigue in the absence of training does not impair learning on a subsequent day

To ensure that the results from the previous experiments were not due to prolonged physical manifestations of fatigue on Day 2, we fatigued a new group of participants on Day 1 but did not expose them to the pinch force task until the second day (FTG_SKILL-DAY2_, *N* =5). On Day 2, this group showed similar learning rates when compared to the control group on Day 1 (mean slope FTG_SKILL-DAY2_ 0.049; p =0.50; see Supplemental Information).

## DISCUSSION

We investigated the effects of fatigue on motor skill learning while excluding performance confounders. As expected, muscle fatigue resulted in lower levels of performance immediately after. However, training under conditions of fatigue in day 1 also affected subsequent training, even in the absence of fatigue. This detrimental effect persisted for almost two complete additional training sessions before performance arrived at equal levels at the end of Day 3. The effect of fatigue on learning was also found on the opposite, not fatigued hand and reverted by a M1 rTMS protocol known to interfere with memory retention (Cantarero et al., 2013a). Further, the detrimental effects of fatigue on skill learning were not present when muscle fatigue was induced but not combined with training. Our results provide evidence for a centrally-mediated deleterious effect of fatigue on motor skill learning beyond impairment in execution.

The observed differences in task performance were mostly driven by lower accuracy rates in the fatigue groups, while movement times were similar across groups. Interestingly, the reduced accuracy was due to a larger number of overshooting errors, which required higher force despite the presence of muscle fatigue. This seemingly counterintuitive finding may be due to sensory attenuation. Sensory attenuation can be defined as the precision with which sensory input from self-generated movement is perceived (Shergill, Bays, Frith, & Wolpert, 2003). Under normal circumstances there is a degree of sensory attenuation associated to voluntary muscle contraction, with lesser attenuation seen for higher force levels (Walsh et al 2011). However, sustained isometric contraction similar to our fatiguing task has been described to directly affect the activity of primary muscle spindle afferents as a consequence of thixotropic properties of intrafusal muscle fibres (Luu, Day, Cole, & Fitzpatrick, 2011). As a result, an underestimation of the absolute applied forces (especially in the lower force range) based on the attenuation of sensory consequences can be expected (Luu et al., 2011, Brooks et al., 2013). A common experience of this effect in everyday life is the Kohnstamm’s phenomenon, where preconditioning of the muscle spindles with isometric contraction has significant effect on position sense and sense of effort (Hagbarth and Nordin, 1998. Therefore, it is conceivable that changes in sensory attenuation induced by fatigue led to impaired force control on Day 1.

Alternatively, it has recently been suggested that fatigue might distort visual perception. This has been shown from both egocentric perspective (Witt and Profitt 2003) and allocentric perspective (Kuppuswamy et. al., 2016). For example, participants who experienced high-level physical exertion systematically overestimated the length of a visually presented line, which could result in target overshooting (Kuppuswamy et al 2016).

How these processes exert their effect on subsequent training sessions and take multiple days to wash out is not clear. Yet, in our understanding, the observed result cannot simply be explained by context specificity. It has been argued that, because fatigue leads to changes in the pattern and intensity of muscle activation as well as to changes in sensory feedback, what has been learned under fatigue can only show limited transfer to performance in the unfatigued state and vice versa (Barnett, Ross, Schmidt, & Todd, 1973; Janet L, 2012). While limited transfer can explain an initially lower performance of the fatigued group on Day 2, it cannot account for the lower learning rates throughout Day 2 and sustained effect on performance up to Day 3.

We propose that subsequent training is impaired because specific central learning mechanisms are affected under fatigue. For instance, it is conceivable that aspects of task performance under fatigue (e.g. changes in the pattern and intensity of muscle activation) are remembered and recalled during subsequent practice even though the system properties have changed between Day 1 and Day 2. Muscular fatigue induces peripheral as well as central changes, such as decreases of motor cortical drive to the muscles followed by depression of motor cortex excitability (Gandevia, Allen, & McKenzie, 1995a; J. L. Taylor, Butler, & Gandevia, 2000; Zanette et al., 1995). This is particularly interesting as motor cortex function has been attributed an essential role for skill learning and retention (Kawai et al., 2015; Shmuelof & Krakauer, 2011). Supporting evidence to central mechanisms also comes from Takahashi and colleagues who found that subjects produced higher forces during a second exposure to a force field if they had practiced the task under fatigue compared to practice in a non-fatigued condition (Takahashi, 2006). A central mechanism is also indicated by our experiment 3 results. Here, we found that participants who were fatigued on Day 1, but received disruptive rTMS over the motor cortex (known to affect retention (Cantarero et al., 2013a) after training expressed higher learning rates than participants who only received rTMS over a control cortical site on the subsequent day. Indeed, after rTMS no differences in learning rate could be found when compared to non-fatigued controls on day 2. Therefore, our results indicate that learning under fatigue can lead to the formation of specific memories that are not helpful to subsequent training in a non-fatigued state, slowing down overall learning.

The persistent limited skill acquisition following training under fatigue may be attributed to time-dependent differences in the contribution of explicit and automatized memory-based processes (J. A. Taylor & Ivry, 2012). Explicit strategic processes generally have been associated with early stages of learning, when the difference between the movement goal and the chosen motor command is large (J. A. Taylor & Ivry, 2012). In this stage, exploration of the manifold is believed to lead towards selection of the optimal (or close to optimal) solution strategy. This is then followed by a gradual refinement of the chosen action sequence and smaller changes in behaviour. Thus, it is conceivable that aspects of task performance under fatigue are indeed retained and perceived as the optimal movement strategy on the second day, leading to slower performance gains. In other words, participants that learned the “optimal motor commands for the fatigued state” may lack a de novo exploratory stage in the subsequent exposures to the task, resulting in continued use of a strategy that is suboptimal for learning which, in turn, results in lower learning rates.

## CONCLUSION

We tested motor learning of a skill task under conditions of fatigue. We found that learning in a fatigued state results in detrimental effects on overall task acquisition. These phenomenon is present above and beyond the deleterious consequences of fatigue on performance and appears to be, at least in part, centrally mediated. These observations need to be carefully considered when designing training protocols such as in sports or musical performance as well as for rehabilitation programs. While conditions of fatigue during sports or performing arts can occur by chance or by overachieving attitudes, rehabilitation programs are particularly at risk because patients with neurological conditions such as those following stroke or multiple sclerosis frequently experience fatigue.

## METHODS

### Participants

A total of 103 healthy participants were recruited from two centres (Johns Hopkins University and University College London). None of the participants suffered from any neurological or psychiatric disorder, nor were they taking any centrally-acting prescribed medication. The experiments were approved by the respective ethics boards at Johns Hopkins School of Medicine Institutional Review Board and the North West London Research Ethics Committee in accordance to the Declaration of Helsinki, and written informed consent was obtained from all participants. For the first experiment sample size was chosen in line with previously reported effect sizes in motor skill learning studies (Cantarero et al., 2013a; Cantarero, Tang, O’Malley, Salas, & Celnik, 2013b; Reis et al., 2009).

### Motor task

For each experiment, participants were seated in front of a computer monitor and given a force transducer to hold between the thumb and index fingers of their dominant hand. During each trial, participants were instructed to produce isometric pinch presses at different force levels to control the motion of a cursor displayed on the screen. Increasing force resulted in the cursor moving horizontally to the right. Participants were instructed to increase and decrease their pinching force to navigate the cursor through the following sequence: start-gate1-start-gate2-start-gate3-start-gate4-start-end; see Fig 6. The cursor movement followed a logarithmic transduction of the applied pinch force as described in previous studies (Reis et al., 2009). This task has been widely used to study skill learning (e.g., (Reis et al., 2009, Cantarero et al., 2013). It involves two components of learning, speed and accuracy, which we could explore independently (see also Data Analysis and Supplemental Information).

**Figure 6.**
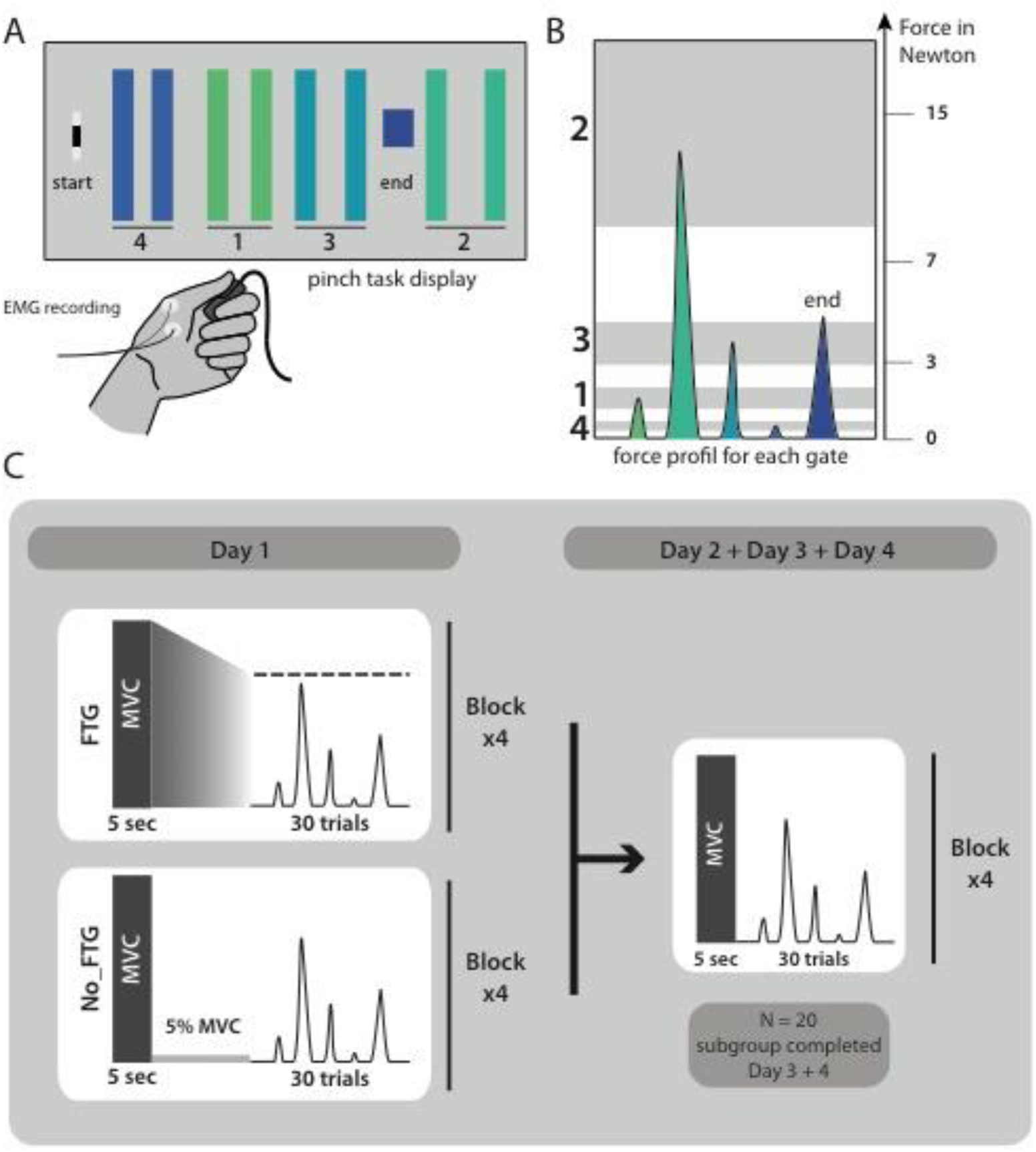
Pinch force skill task and study design for experiment 1.

### Design

#### Experiment 1: Determining the effect of fatigue on temporal aspects of motor skill learning

38 participants (23 women, mean age 22.2, ±2 years, all right-handed) were recruited and randomly assigned to one of two groups, a fatigue group (FTG, *N* =20) and a non-fatigue group (NoFTG, *N* =18). All participants underwent ∼45 min testing sessions on two or four consecutive days; see Fig. 6. Sessions took place between 9 a.m. and 6 p.m. and were separated by 24h (±1 hour). Morning and afternoon sessions were counterbalanced between groups and subjects performed both sessions at similar times. Each day, both groups performed the isometric pinching task (see motor task) for four blocks of 30 trials each. At the start and the end of each experimentation sessions, participants in both groups were asked to press the force transducer with their maximum force for 5 seconds in order to assess the maximum voluntary contraction (MVC). On Day 1, the FTG was instructed to sustain MVC until the produced force dropped to the level of the upper limit for gate 2, the target that requires the largest force production. Thus, the induction of fatigue always stayed above the force level needed for task execution. Time to fatigue was 68.91, SD 32.23 seconds on average. To counterbalance the amount of time of the fatigue induction, the non-fatigue group was asked to sustain 5% of their MVC over a matched period of time. On Day 2, the design was identical for both groups: assessment of MVC at the start and end of the session, with four blocks of skill task without break in between. To determine how long fatigue influenced motor skill learning, a subgroup (*N* =24, FTG and NoFTG both *N* =12) participated in two extra days of experimentation. The design in Days 3 and 4 was identical to Day 2 for both groups.

#### Experiment 2: Determining effects of fatigue on motor skill learning measured by transfer of skill

To investigate motor skill learning within one session while avoiding the execution confounder, we assessed the amount of inter-manual skill transfer in the unfatigued, untrained left hand after participants trained with the right hand with or without fatigue. 20 participants were recruited to take part in experiment 2 (17 women, mean age 20.1 ±3.0 years, all right-handed). Participants were again randomly assigned to either a fatigued or a non-fatigued group (FTG_TRANSFER_, *N* =10; NoFTG_TRANSFER_, *N* =10) and tested in a single session. The task design was identical to the first day in experiment 1. Additionally, at the beginning and end of the session, participants completed one block of 15 trials with their unfatigued left hand; for visualisation of the study design see Supplemental Information.

#### Experiment 3: Testing whether effects of fatigue on learning are centrally mediated

In experiment 3, a new group of 40 healthy participants were recruited (15 women, mean age 23.5 ±2.9 years, 4 left-handed, one ambidextrous) and randomly assigned to one of three groups: a non-fatigue group (NoFTG_SHAM_, *N* =10), or one of two Fatigue groups (FTG_SHAM_, *N* =15 or FTG_M1_, *N* =15). The overall study design for all three groups was similar to experiment 1; for visualization of the study design see Supplemental Information. In addition, at the end of Day 1, participants received depotentiation TMS (DePo) either over their registered “M1 hot-spot” (FTG_M1_; see also TMS section) or a control location (FTG_SHAM_ & NoFTG_SHAM_). The depotentiation stimulation is a shorter form of continuous theta burst stimulation with TMS which has been shown to reverse potentiating plasticity (Huang et al., 2010; Huang, Edwards, Rounis, Bhatia, & Rothwell, 2005; Huang, Rothwell, Edwards, & Chen, 2008) and disrupt skill retention (Cantarero et al., 2013a). DePo stimulation was administered in a double-blind fashion. A researcher not involved in the behavioral portion of the study delivered the stimulation, while those conducting the behavioral training were blinded to the stimulation location and protocol.

### TMS

Transcranial magnetic stimulation (TMS) was administered with a figure-eight coil (wing diameter =70 mm) connected to a Magstim 200 stimulator (Magstim, UK). Using TMS, We located the “hot-spot” of the abductor pollicis brevis muscle in the task-relevant hand at rest according to standardized procedures (Chen et al., 2008; Rossini et al., 1994). The stimulus intensity that elicited a motor evoked potential (MEP) with a peak-to-peak amplitude of approximately 1 mV was established (Stimulus intensity 1mV, S1mV) to assess corticomotor excitability. Then 18 MEPs were recorded using the same intensity before the task, directly after the task, and after depotentiation on Day 1 as well as before and after the task at Day 2. The parameters for depotentiation, were based on previous reports (Cantarero et al., 2013a; Huang et al., 2010), consisted of bursts of three pulses at 50 Hz repeated at 200 ms intervals at an intensity of 70% of the rMT for 10 seconds. Using the 10-20 electroencephalogram coordinate system, Pz was used as a control stimulation location (SHAM stimulation). Stereotactic neuronavigation (BrainSight, Rogue Research, Montreal, Quebec, Canada) was used to track coil position within sessions. EMG activity from the abductor pollicis brevis muscle was recorded using surface electrodes taped in a belly-tendon orientation. Data was recorded with an AMT-8 (Bortec Biomedical Ltd; sampling rate 5000 Hz, amplification 1000x, band-pass filtering 10-1000 Hz) and saved for offline analysis.

### Data analysis

For analysis of MVC see Supplemental Information.

### Analysing Movement Time/Error Rate

For each trial in the skill task, movement time and error rate were recorded: movement time was defined as the duration from movement onset (forced controlled cursor leaving start position) to reaching the end gate. Error rate was defined as the percentage of trials per block in which participants under- or overshot at least one of the five targets.

### All experiments

Movement time and error rate were compared using rmANOVA with the within-subject factor ‘block’ (four levels: b1, b2, b3 and b4) and the between-subject factor ‘group’ (Exp. 1 & 2: two levels, Exp. 3: three levels) for each single day (see Supplemental Information).

### Analysing motor skill

To quantify motor performance, we calculated a skill measure, composed of movement time and error rate. As done in prior studies, the skill measure was calculated as: a=(1-error rate)/ [error rate (ln(movement time)b)], where b is 5.424 as predefined for this particular task in prior studies (Reis et al., 2009, Cantarero et al., 2013(Mawase, Uehara, Bastian, & Celnik, 2017; Spampinato & Celnik, 2017)).

To study learning (rate of change in performance) during the motor task, we plotted the number of blocks on the x-axis and the skill measured on the y-axis. As the relationship was roughly linear, we fit a robust linear regression model (i.e., f(x) =c*x + b) for each group; the robust function disregards outliers by estimating an iteratively reweighted least square algorithm. This provided a more parsimonious model than ANOVA, providing a single easily interpretable measure of learning rate given by the slope c. We were particularly interested in measuring and comparing the learning rates between days and groups. When only two blocks were tested (i.e. the left hand in experiment 2) we took the difference between block2 – block1 as a measure of learning rate.

Differences in learning rates for each experiment were assessed using a permutation testing procedure. Assuming the null hypothesis of no group difference, participants were randomly reassigned to the two groups, and the difference in regression coefficients between the resampled groups was computed. This procedure was repeated 10,000 times, allowing us to generate a null distribution for the difference between regression coefficients assuming no group differences. The proportion of resampled values that exceeded the true observed difference was used to compute p-values and determine statistical significance. Under the null hypothesis, the true difference in learning rates between the two groups should lie within the distribution of these randomly generated differences, with extreme values providing evidence against the null hypothesis.

Prior to application of any parametric tests, the normality of the dependent variables was assessed using Shapiro-Wilk tests and quantile-quantile plots. A log-transformation was applied to correct for any non-normal data. All ANOVA results were Greenhouse-Geisser corrected if the assumption of sphericity was violated. Student’s t-test was used to assess group differences. Results were considered significant at p<0.05, and Bonferroni correction was applied to correct for multiple comparisons. All data are expressed as mean ±standard error unless stated otherwise. Statistical analyses were performed using SPSS 22.0 ® and custom-written MATLAB routines.

## ACKNOWLEDGEMENTS

We thank Claudia Ammann for conducting the depotentiation stimulation.

## SUPPLEMENTAL INFORMATION

### Control experiment

To test possible long lasting effects of fatigue on performance rather than learning, we induced fatigue on Day 1 as previously described, but participants were not trained on the skill task; instead, they took breaks matched to the time the other groups needed to finish one block on average (FTG_SKILL-DAY2_, *N* =5). On Day 2, this group showed similar learning rates when compared to the control group on Day 1 (mean slope FTG_SKILL-DAY2_ 0.049; p =0.50; Fig S1).

### Analysis of MVC

MVC was expressed as the average absolute force applied on the force transducer for 5 seconds. Additionally, we recorded surface EMG from the first dorsal interosseous and the adductor pollicis brevis.

#### Experiment 1

To ensure that the chosen condition was reliably inducing fatigue over the whole duration of the task, we assessed the decrease of MVC on Day 1 using repeated-measures ANOVA (rmANOVA) with the within-subject factor ‘time’ (five levels: preB1, preB2, preB3, preB4, postB4) for the fatigued group. On Day 2, we assessed changes in MVC using an rmANOVA with the within-subject factor ‘time’ (two levels: pre, post) and the between-subject factor group (two levels: NoFTG, FTG).

#### Experiment 2

We compared differences in MVC before and after the experiment, denoted ΔMVC(pre-post), between the two groups (FTG_TRANSFER_ versus NoFTG_TRANSFER_) using a one-way ANOVA.

#### Experiment 3

Differences in Δ MVC(pre-post) between days and groups were determined with rmANOVA with the within-subject factor ‘day’ (two levels: Day 1 and Day 2) and the between-subject factor ‘group’ (three levels: NoFTG_SHAM_, FTG_SHAM_, FTG_M1_).

#### Experiment 4 & 5

MVC before and after the task (ΔMVC(pre-post)) was compared to determine the induction of fatigue using a paired two-tailed Student’s t-test.

### Induction of Fatigue

#### Experiment 1

To evaluate if fatigue was reliably induced over the whole course of the task in the fatigue condition, we performed an rmANOVA with the within-subject factor time (5 levels: preB1, preB2, preB3, preB4, postB4). On day 1, the fatigued group showed a significant drop of MVC over time F(4,76) =6.46, p<0.001. In the NoFTG group paired comparisons between MVC before and after the task showed no significant difference (p =0.16). On day 2, no difference in MVC for the two groups was found (time (F(1,34) =2.90, p =0.10; group (F(1,34) =1.75, p =0.20; time*group (F(1,34) =0.19, p =0.67).

#### Experiment 2

For experiment 2 we found a significant interaction of time*group for the right hand (F(1,18) =4.823, p =0.041). There was a significant difference for ΔMVC_right between the NoFTG_TRANSFER_ and FTG_TRANSFER_ (t(1,18) =−2.2, p =0.041, NoFTG_TRANSFER_ −10 Newton ±7.18 indicating higher MVC for the control group after the task, FTG_TRANSFER_ 7.77 Newton ±3.73 indicating lower forces post task for the fatigue group as expected). No significant interaction of time*group nor an significant difference of Δ MVC_left between groups was found (F(1,18) =0.789, p =0.748; t(1,18) =0.328, p =0.747).

#### Experiment 3

Looking at ΔMVC(pre-post), we found an significant interaction between day*group (F(2,37) =3.390, p =0.044). On Day 1 the ΔMVC of the control group was different from both fatigued groups, while no significant difference was found between them (F(2,37) =9.755, p <0.001, NoFTG vs FTG_SHAM_, t(2,37) =−3.49, p =0.004, vs FTG_M1_, t(2,37) =−4.27, p<0.001, FTG_SHAM_ vs FTG_M1_, t(2,37) =−0.87, p =0.9, NoFTG_SHAM_ −7.32 Newton ±3.64, FTG_SHAM_ 6.9 Newton ±1.97, FTG_M1 10 Newton ±2.81). On Day 2 no significant difference in ΔMVC between all groups was found (F(2,37) =1.631, p =0.209).

#### Experiment 4 & 5

For experiment 4 we found a significantly lower MVC post compared to pre task on both days (Exp. 4 day 1 t(1,4) =2.91, p =0.044, day 2 t(1,4) =6.3, p =0.004). For experiment 5, subjects only completed the fatigue condition without the task on day 1 and MVC significantly dropped pre to post, while no difference was found on Day 2 were subjects only completed the task without the fatigue condition (Exp. 5 day 1 t(1,4) =2.92, p =0.043; day 2 t(1,4) =1.91, p =0.129).

#### Movement Times and Error Rates

The implemented skill measure consists of two components: speed and accuracy of the movement. Because speed and accuracy are linked inversely (lower speed allows for more accuracy and vice versa), comparing accuracy at different speed level can be difficult to interpret(Hardwick, Rajan, Bastian, Krakauer, & Celnik, 2016). We separately looked at the two components of the implemented skill measure to determine if differences between the fatigued and the control group were based on changes in speed, accuracy or both.

#### Experiment 1

Results showed that differences in the skill measure between the fatigued and the control group were based on divergent error rates while movement time stayed similar. Participants got faster from B1 to B4 but there was no significant difference between groups or an interaction between block*groups on both days (Day 1: block: F(3,108) =90.692, p <0.001; group: (F(1,36) =0.019, p =0.892), block*group: (F(3,36) =0.220, p =0.760; Day 2: block: F(3,108) =18.425, p <0.001; group: (F(1,36) =0.569, p =0.456), block*group: (F(3,36) =1.423, p =0.240). Regarding error rate, there was significant effect of ‘group’ (F(1,36) =2.072, p =0.031) and no significant effects of ‘block’ and ‘block*group’ on Day 1 (F(3,108) =8.53, p =0.44; F(3,36) =0.388, p =0.7). On Day 2, we found significant effect of ‘block’ (F(3,108) =3.827, p=0.012) and no significant effect of ‘group’ or ‘block*group’ (F(1,36) =2.155, p =0.151; (F(3,36) =0.511, p =0.675). To be better understand the origin of the lower skill rate of the fatigued participants, we additionally analyzed the applied forces of each group. Surprisingly, the fatigued group exerted overall more force than the control group (t(1,36) =−2.61, p =0.013, NoFTG 6.9 Newton ±0.012, FTG 7.8 Newton ±0.048). Interestingly overshooting was only significantly different between groups for the lower force targets T1 and T4 (t(1,36) =−3.172, p =0.003, respectively t(1,36) =−2.537, p =0.015) but not the higher force targets T2 and T3 (t(1,36) =−1.036, p =0.307, respectively t(1,36) =−2.104, p =0.051).

#### Experiment 2

Movement time was similar between both groups for the right and the left hand (Right hand: F(1,18) =1.975, p =0.177; Left hand: F(1,18) =0.392, p =0.539), while error rates were significantly different (Right hand: F(1,18) =26.075, p <0.001; Left hand: F(1,18) =29.555, p <0.001).

#### Experiment 3

As expected, in experiment 3 we found differences in error rate but not movement time between groups (movement time/Day 1: F(2,37) =0.817, p =0.449; /day 2: F(2,37) =0.075; error rate/day 1: F(2,37) =9.358, p =0.001; day 2: F(2,37) =4.995, p =0.012). There was no interaction between block*group, F(6,37) =0.175, p =0.983).

## Study Design

### Visualization of study designs

#### Experiment 2

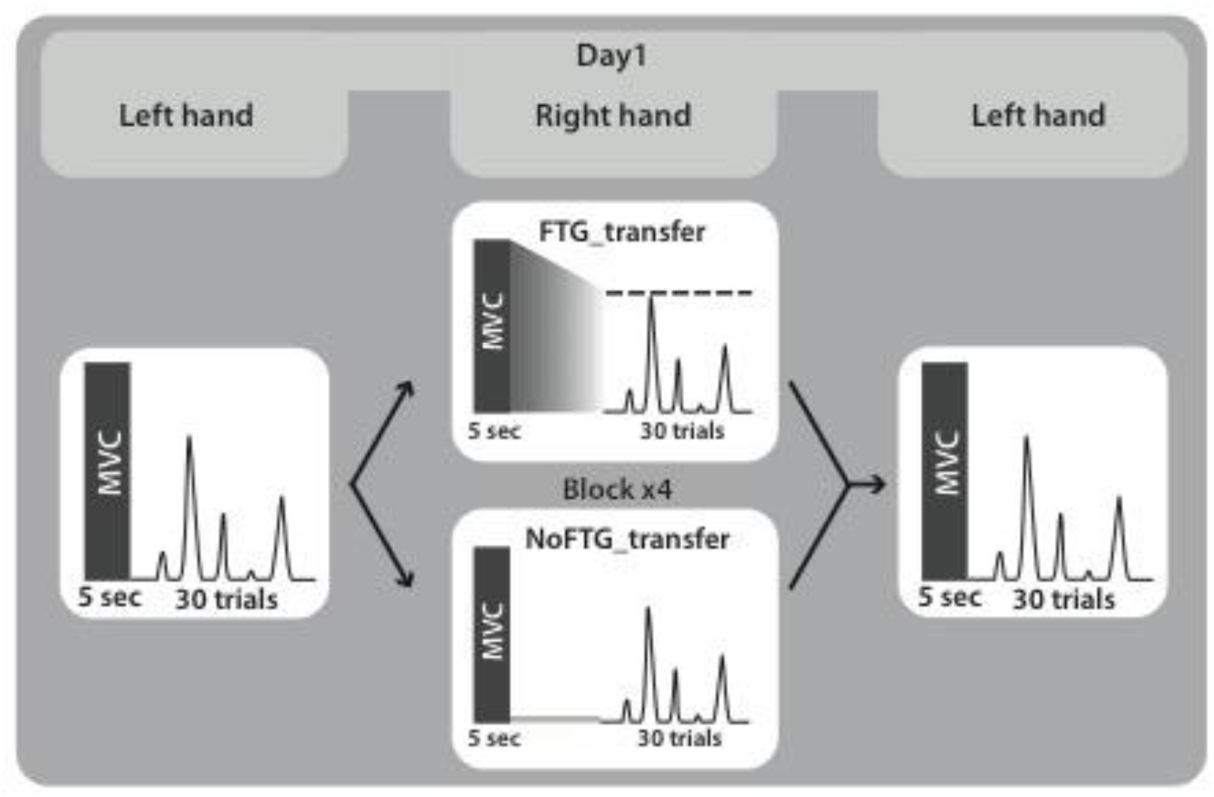

#### Experiment 3

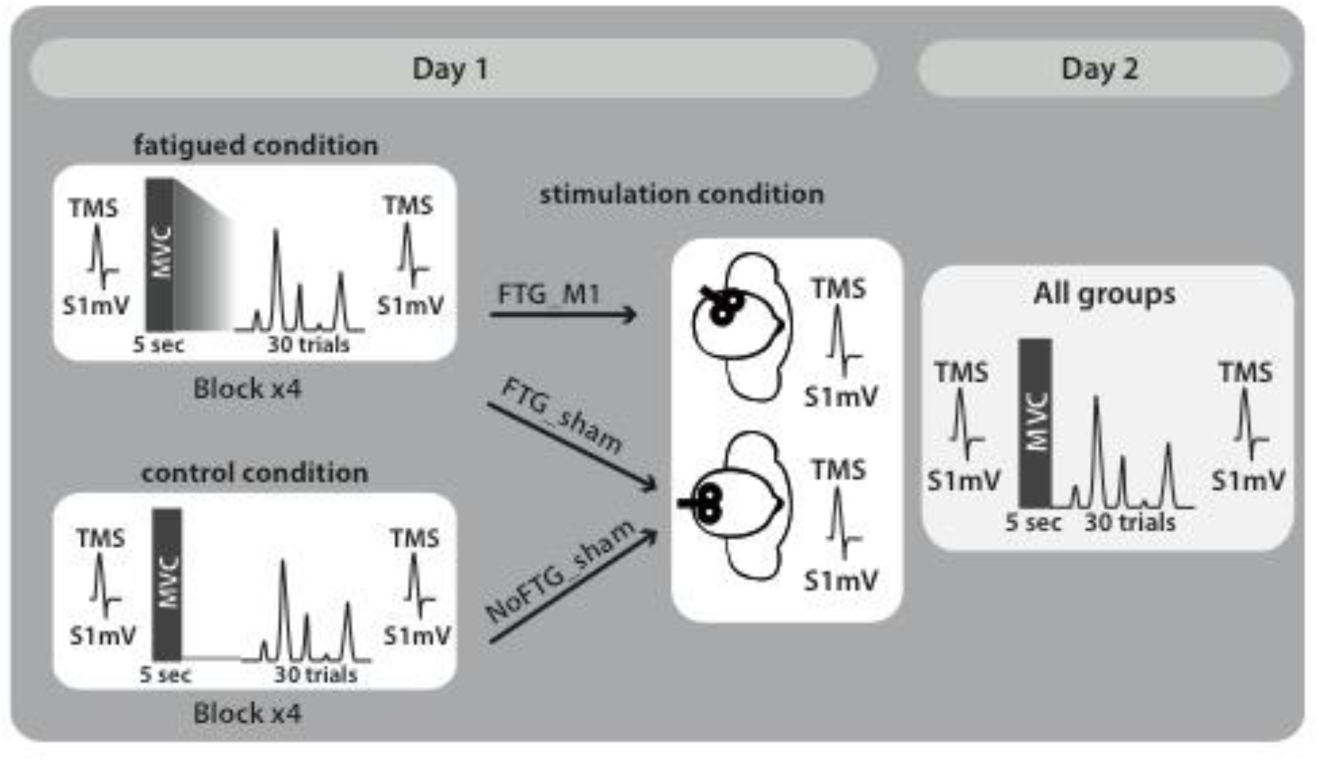

---

**Figure S1:**
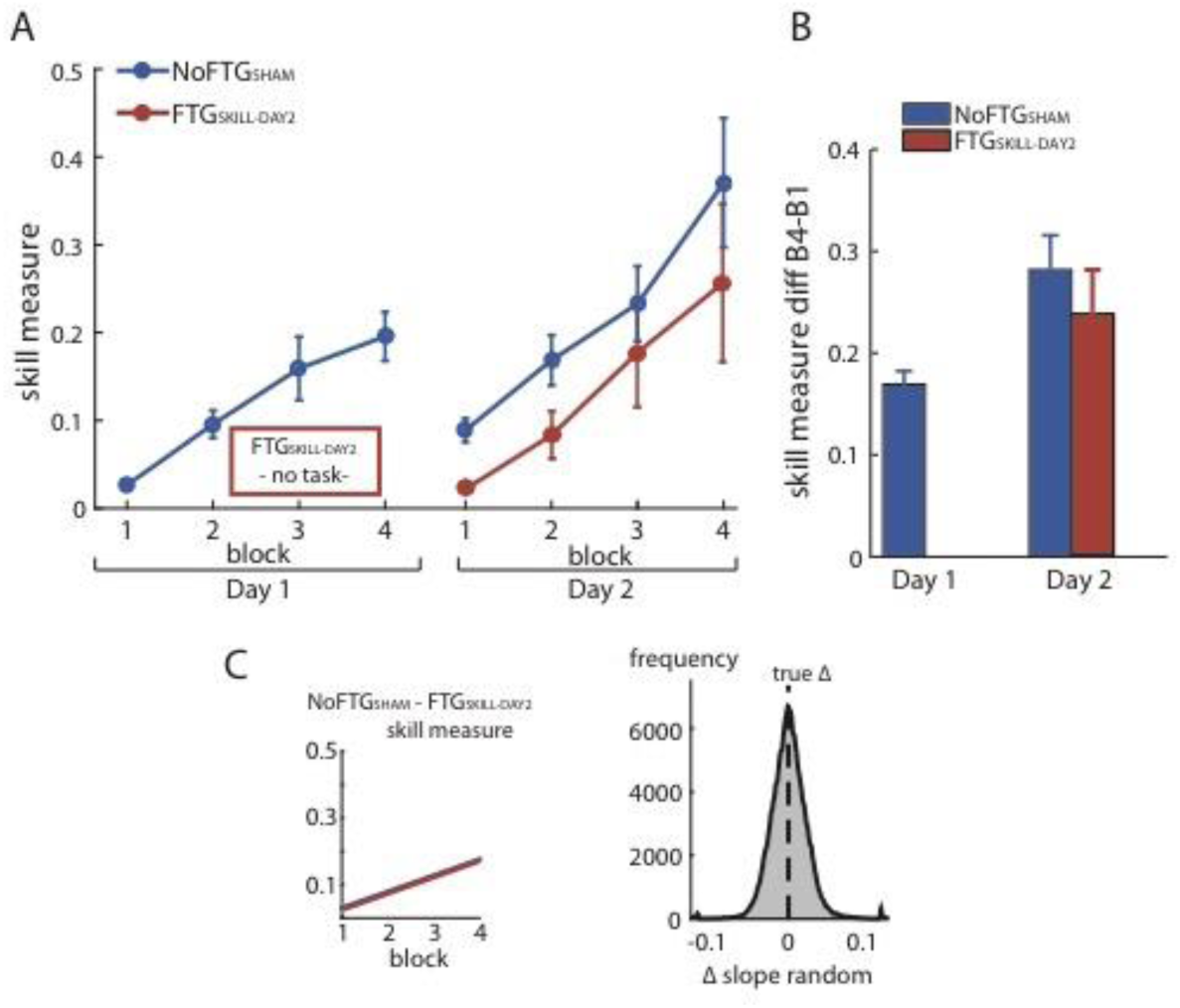
Fatiguing without training did not affect learning on Day 2. Panel (A) shows changes in skill measure over the course of four blocks on two consecutive days for the non-fatigued group (NoFTG_SHAM_, same control group as in experiment 3) and on Day 2 for the FTG_SKILL-DAY2_ (participants were fatigued on Day 1, but did not learn the task until Day 2 when they trained in the absence of fatigue). FTG_SKILL-DAY2_ learned the task on Day 2 to a similar extent as the non-fatigued group on Day 1. As expected, the FTG_SKILL-DAY2_ performed at a lower level compared to the non-fatigued group on Day 2. Panel (B) shows the difference in performance between block 4 to block 1 for both groups across days, the relative improvement in the FTG_SKILL-DAY2_ was comparable to the control group on both days. Panel (C) shows the learning rates for the control group on Day 1 compared to the FTG_SKILL-DAY2_. No differences in learning rates were found using permutation testing.

**Table 1:**
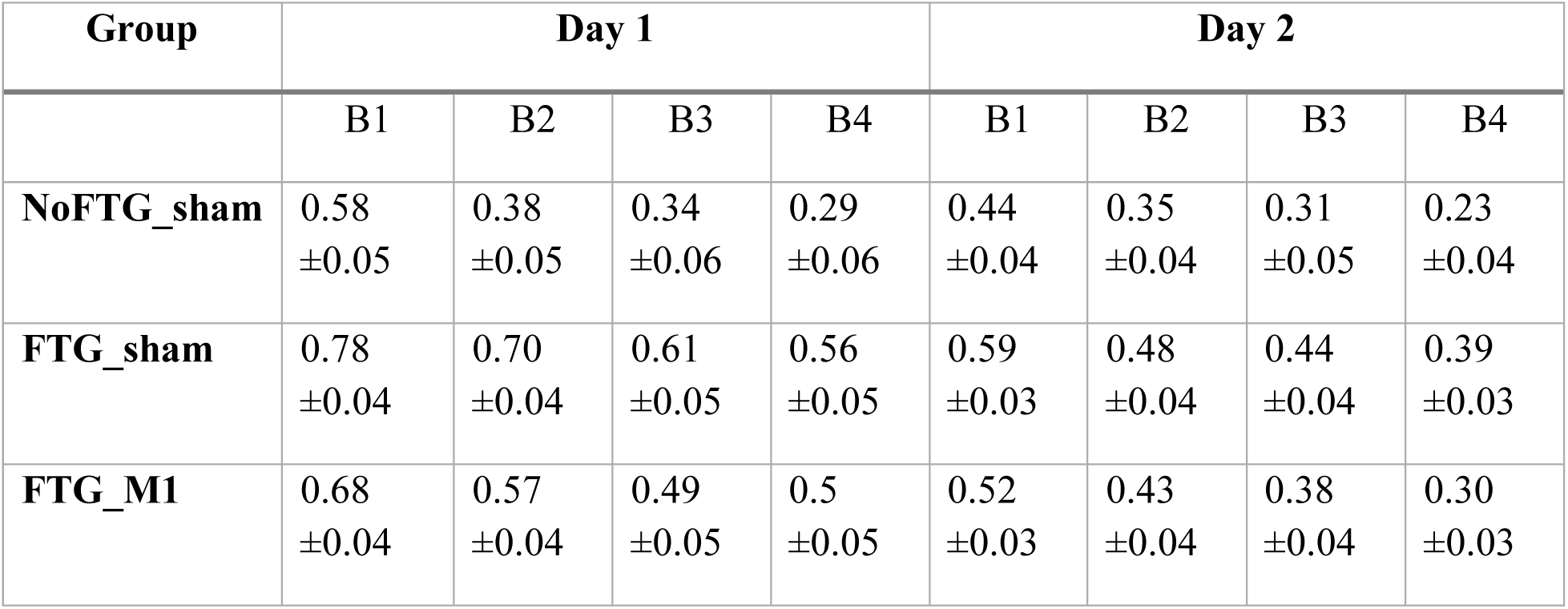
error rate experiment 3.

## REFERENCES

Barnett, M. L., Ross, D., Schmidt, R. A., & Todd, B. (1973). Motor skills learning and the specificity of training principle. Research Quarterly, 44(4), 440–447.

Boyas, S., & Guével, A. (2011). Neuromuscular fatigue in healthy muscle: Underlying factors and adaptation mechanisms. Annals of Physical and Rehabilitation Medicine, 54(2), 88–108. http://doi.org/10.1016/j.rehab.2011.01.001

Cahill, L., McGaugh, J. L., & Weinberger, N. M. (2001). The neurobiology of learning and memory: some reminders to remember. Trends in Neurosciences, 24(10), 578–581.

Camus, M., Ragert, P., Vandermeeren, Y., & Cohen, L. G. (2009). Mechanisms controlling motor output to a transfer hand after learning a sequential pinch force skill with the opposite hand. Clinical Neurophysiology: Official Journal of the International Federation of Clinical Neurophysiology, 120(10), 1859–1865. http://doi.org/10.1016/j.clinph.2009.08.013

Cantarero, G., Lloyd, A., & Celnik, P. (2013a). Reversal of Long-Term Potentiation-Like Plasticity Processes after Motor Learning Disrupts Skill Retention. Journal of Neuroscience, 33(31), 12862–12869. http://doi.org/10.1523/JNEUROSCI.1399-13.2013

Cantarero, G., Tang, B., O’Malley, R., Salas, R., & Celnik, P. (2013b). Motor learning interference is proportional to occlusion of LTP-like plasticity. Journal of Neuroscience, 33(11), 4634–4641. http://doi.org/10.1523/JNEUROSCI.4706-12.2013

Galea, J. M., Vazquez, A., Pasricha, N., de Xivry, J.-J. O., & Celnik, P. (2011). Dissociating the roles of the cerebellum and motor cortex during adaptive learning: the motor cortex retains what the cerebellum learns. Cerebral Cortex (New York, N.Y.: 1991), 21(8), 1761–1770. http://doi.org/10.1093/cercor/bhq246

Gandevia, S. C., Allen, G. M., & McKenzie, D. K. (1995a). Central fatigue. Critical issues, quantification and practical implications. Advances in Experimental Medicine and Biology, 384, 281–294.

Gandevia, S. C., Enoka, R. M., McComas, A. J., Stuart, D. G., & Thomas, C. K. (1995b). Neurobiology of Muscle Fatigue. In Fatigue (Vol. 384, pp. 515–525). Boston, MA: Springer US. http://doi.org/10.1007/978-1-4899-1016-5_39

Hardwick, R. M., Rajan, V. A., Bastian, A. J., Krakauer, J. W., & Celnik, P. A. (2016). Motor Learning in Stroke. Neurorehabilitation and Neural Repair. http://doi.org/10.1177/1545968316675432

Huang, Y.-Z., Edwards, M. J., Rounis, E., Bhatia, K. P., & Rothwell, J. C. (2005). Theta burst stimulation of the human motor cortex. Neuron, 45(2), 201–206. http://doi.org/10.1016/j.neuron.2004.12.033

Huang, Y.-Z., Rothwell, J. C., Edwards, M. J., & Chen, R.-S. (2008). Effect of physiological activity on an NMDA-dependent form of cortical plasticity in human. Cerebral Cortex (New York, N.Y.: 1991), 18(3), 563–570. http://doi.org/10.1093/cercor/bhm087

Huang, Y.-Z., Rothwell, J. C., Lu, C.-S., Chuang, W.-L., Lin, W.-Y., & Chen, R.-S. (2010). Reversal of plasticity-like effects in the human motor cortex. The Journal of Physiology, 588(Pt 19), 3683–3693. http://doi.org/10.1113/jphysiol.2010.191361

Janet L T. (2012). Motor Control and Motor Learning Under Fatigue Conditions. In Routledge Handbook of Motor Control and Motor Learning. Routledge. http://doi.org/10.4324/9780203132746.ch17

Kantak, S. S., & Winstein, C. J. (2012). Learning-performance distinction and memory processes for motor skills: a focused review and perspective. Behavioural Brain Research, 228(1), 219–231. http://doi.org/10.1016/j.bbr.2011.11.028

Kawai, R., Markman, T., Poddar, R., Ko, R., Fantana, A. L., Dhawale, A. K., et al. (2015). Motor cortex is required for learning but not for executing a motor skill. Neuron, 86(3), 800–812. http://doi.org/10.1016/j.neuron.2015.03.024

Kuppuswamy, A. (2017). The fatigue conundrum. Brain: a Journal of Neurology, 140(8), 2240–2245. http://doi.org/10.1093/brain/awx153

Kuppuswamy, A., Clark, E. V., Turner, I. F., Rothwell, J. C., & Ward, N. S. (2015). Post-stroke fatigue: a deficit in corticomotor excitability? Brain: a Journal of Neurology, 138(1), 136–148. http://doi.org/10.1093/brain/awu306

Luu, B. L., Day, B. L., Cole, J. D., & Fitzpatrick, R. C. (2011). The fusimotor and reafferent origin of the sense of force and weight. The Journal of Physiology, 589(Pt 13), 3135–3147. http://doi.org/10.1113/jphysiol.2011.208447

Mawase, F., Uehara, S., Bastian, A. J., & Celnik, P. (2017). Motor Learning Enhances Use-Dependent Plasticity. The Journal of Neuroscience: the Official Journal of the Society for Neuroscience, 37(10), 2673–2685. http://doi.org/10.1523/JNEUROSCI.3303-16.2017

Muellbacher, W., Ziemann, U., Wissel, J., Dang, N., Kofler, M., Facchini, S., et al. (2002). Early consolidation in human primary motor cortex. Nature, 415(6872), 640–644. http://doi.org/10.1038/nature712

Perez, M. A., Wise, S. P., Willingham, D. T., & Cohen, L. G. (2007). Neurophysiological mechanisms involved in transfer of procedural knowledge. The Journal of Neuroscience: the Official Journal of the Society for Neuroscience, 27(5), 1045–1053. http://doi.org/10.1523/JNEUROSCI.4128-06.2007

Reis, J., Schambra, H. M., Cohen, L. G., Buch, E. R., Fritsch, B., Zarahn, E., et al. (2009). Noninvasive cortical stimulation enhances motor skill acquisition over multiple days through an effect on consolidation. Proceedings of the National Academy of Sciences of the United States of America, 106(5), 1590–1595. http://doi.org/10.1073/pnas.0805413106

Richardson, A. G., Overduin, S. A., Valero-Cabré, A., Padoa-Schioppa, C., Pascual-Leone, A., Bizzi, E., & Press, D. Z. (2006). Disruption of primary motor cortex before learning impairs memory of movement dynamics. The Journal of Neuroscience: the Official Journal of the Society for Neuroscience, 26(48), 12466–12470. http://doi.org/10.1523/JNEUROSCI.1139-06.2006

Shergill, S. S., Bays, P. M., Frith, C. D., & Wolpert, D. M. (2003). Two Eyes for an Eye: The Neuroscience of Force Escalation. Science, 301(5630), 187–187. http://doi.org/10.1126/science.1085327

Shmuelof, L., & Krakauer, J. W. (2011). Are We Ready for a Natural History of Motor Learning? Neuron, 72(3), 469–476. http://doi.org/10.1016/j.neuron.2011.10.017

Spampinato, D., & Celnik, P. (2017). Temporal dynamics of cerebellar and motor cortex physiological processes during motor skill learning. Scientific Reports, 7, 40715. http://doi.org/10.1038/srep40715

Takahashi, C. D. (2006). Effect of muscle fatigue on internal model formation and retention during reaching with the arm. Journal of Applied Physiology, 100(2), 695–706. http://doi.org/10.1152/japplphysiol.00140.2005

Taylor, J. A., & Ivry, R. B. (2012). The role of strategies in motor learning. Annals of the New York Academy of Sciences, 1251(1), 1–12. http://doi.org/10.1111/j.1749-6632.2011.06430.x

Taylor, J. L., Butler, J. E., & Gandevia, S. C. (2000). Changes in muscle afferents, motoneurons and motor drive during muscle fatigue. European Journal of Applied Physiology, 83(2-3), 106–115. http://doi.org/10.1007/s004210000269

Tolman, E. C., & Honzik, C. H. (1930). “Insight” in Rats.

Zanette, G., Bonato, C., Polo, A., Tinazzi, M., Manganotti, P., & Fiaschi, A. (1995). Long-lasting depression of motor-evoked potentials to transcranial magnetic stimulation following exercise. Experimental Brain Research. Experimentelle Hirnforschung. Expérimentation Cérébrale, 107(1), 80–86.

